# Stomata, a vulnerability in the plant defence against phytophagous mites that ABA can overcome

**DOI:** 10.1101/2023.09.29.555308

**Authors:** Irene Rosa-Diaz, James Rowe, Ana Cayuela-Lopez, Vicent Arbona, Isabel Diaz, Alexander M. Jones

## Abstract

Arthropod herbivory possess a significant threat to crop yield, prompting plants to employ intricate defense mechanisms against pest feeding. The generalist pest, *Tetranychus urticae*, inflicts rapid damage and remains a challenge due to its broad target range. In this study, we explored *Arabidopsis thaliana’s* response to *T. urticae* infestation, revealing the induction of abscisic acid (ABA), a hormone typically associated with abiotic stress adaptation, including stomatal closure during water stress. Leveraging a FRET-based ABA biosensor (nlsABACUS2-400n), we observed elevated ABA levels in various leaf cell types post-mite feeding. While ABA’s role in pest resistance or susceptibility has been debated, an ABA-deficient mutant exhibited increased mite infestation, alongside intact canonical biotic stress signalling, indicating an independent function of ABA in mite defense. Through genetic and pharmacological interventions targeting ABA levels, ABA signalling, stomatal aperture, and density, we established that ABA-triggered stomatal closure effectively hinders mite feeding and minimizes leaf cell damage. This study underscores the critical interplay between biotic and abiotic stresses in plants, highlighting how the vulnerability to mite infestation arising from open stomata, crucial for transpiration and photosynthesis, underscores the intricate relationship between these two stress types.

## Introduction

Plants and phytophagous arthropods have coevolved in an ‘arms race’ of mutual antagonism between herbivory strategies and plant defences. Successful feeding by insect and acari pests on compatible crop plants causes substantial yield losses and represents a threat to global food security. Successful plant defence starts in part with the molecular recognition of Herbivore Associated Molecular Patterns (HAMPs) and Damage-Associated Molecular Patterns (DAMPs). Through these early warning cues, plants prompt signal transduction pathways and transcriptional reprogramming that result in the production of chemical and physical defences (Stahl et al., 2018; Garcia et al., 2021). This cascade of events requires fine-tuned control mediated by multiple regulatory factors to induce defence regimes appropriate to herbivore species, plant host and environment conditions. Common chemical responses to herbivore damage include the accumulation of reactive oxygen species (ROS), calcium, and several hormones considered to be core modulators of immune signalling, i.e. jasmonic acid (JA), jasmonoyl-isoleucine (JA-Ile) and salicylic acid (SA), (Erb & Reymond, 2019). These responses are also shared with many plant-pathogen interactions (Bari and Jones, 2009). Physical defences against pests vary considerably, for example lignified and suberised root and stem barriers, aerial trichomes and epidermal cuticles can all restrict access for feeding (War et al., 2012; Holbein et al., 2019). Induced closure of stomata, as natural entry sites to the leaf interior, has received considerable attention for foliar pathogen responses (i.e. stomatal defence, Melotto *et al*., 2008, 2017). On the other hand, pathogen induced closure of stomata can also contribute to water soaking and microbial pathogenesis later in infection (Hu et al., 2022; Lajeunesse et al., 2023). Stomatal closure has also been observed in response to phytophagous arthropods (Sances et al., 1979; Pincebourde and Casas, 2006; Schmidt et al., 2009), though whether closure constituted a defence mechanism or contributed to the infestation remained unclear.

Abscisic acid (ABA) is the key hormone regulating stomatal aperture, which gates gas exchange for transpiration, photosynthesis and responses to abiotic stimuli (Kuromori et al., 2018a) and is accumulated in response to a number of environmental stimuli including drought, salt and wounding (Christmann et al., 2006). Both chewing and piercing-sucking arthropod pests provoke loss of water, an increase in transpiration and a reduction in stomata conductance. All these phenomena are associated with water stress induced accumulation of ABA and subsequent stomatal closure (Lim et al., 2015). ABA-deficient mutants are more susceptible to chewing herbivory by *Spodoptera exigua* (Thaler and Bostock, 2004) and *Spodoptera littoralis* caterpillars (Bodenhausen and Reymond, 2007), suggesting ABA signalling and potentially stomatal closure could contribute to plant defence. Open stomata are natural apertures where piercing-sucking herbivores like aphids and mites insert their stylets, specialized feeding structures, to access nutrient rich subepidermal cells. Although stylets can also be inserted between epidermal cells, feeding via opening stomata is potentially faster and avoids the damage of cell walls, which can trigger rapid plant defences (Mathen et al., 1988; Bensoussan et al., 2016). In some cases, stomata are even preferred places to lay herbivore eggs (DeClerck and Steeves, 1988). On the other hand, the caterpillar *Helicoverpa zea* was found to actively induce stomatal closure to reduce release of herbivore-induced plant volatiles that can recruit *H. vea* predators (Lin et al., 2021). More direct regulation of ABA by piercing-sucking herbivores is also possible as the saliva of the aphid *Myzus persicae* up-regulated genes that also responded to ABA treatment (Hillwig et al., 2016). In this interaction, ABA contributes to susceptibility to aphid infestation, indicating that the role of ABA signalling in herbivory defence could also be negative.

ABA is perceived by PYRABACTIN RESISTANCE (PYR)/PYR-like (PYL) or REGULATORY COMPONENT OF ABA RECEPTOR (RCAR) family receptor proteins (Gonzalez-Guzman et al., 2012). The accumulation of ABA in stomatal guard cells, for example during water stress, is transduced via PYR/PYL/RCAR receptors to a complex signalling network (Ma et al., 2009). Core components include inhibitory PROTEIN PHOSPHATASE 2C (PP2C)-type proteins as well as positive regulators of ABA signalling including, SUCROSE NON-FERMENTING 1-RELATED PROTEIN KINASE 2 (SnRK2)-type protein kinases as well as ABRE-BINDING FACTOR (ABF) and ABA Insensitive (ABI)-type transcription factors (Hsu et al., 2021). ABA effects comprise changes in redox homeostasis, guard cell-specific kinase regulation, and changes in cytosolic Ca^2+^ levels. These events lead to stomatal closure while also regulating numerous non-stomatal functions (Neill et al., 2008; Dinh et al., 2013; Ng et al., 2014). ABA regulation of these components also occurs in many plant-pathogen interactions, and as for plant-herbivore interactions, ABA has been found to both promote and antagonize microbial defences (Bari and Jones, 2009; Lim et al., 2015)

One explanation for contrasting roles for ABA in diverse plant defence responses is ABA crosstalk with ROS, calcium, and defence hormone signalling, for example through inhibition of SA signalling (De Torres-Zabala et al., 2007; Ton et al., 2009). ABA activation of primed JA-regulated defence responses in *Arabidopsis thaliana* contributes to induced resistance against *Pieris rapae* herbivory (Vos et al. 2013). ABA can also regulate JA responses via direct interaction of PYL5/6 ABA receptors with MYC transcription factors (Aleman et al., 2016) and ABA amplifies JA-dependent defence responses as part of the signal transduction pathway in plants elicited by oral secretions of insects (Dinh et al., 2013). Thus, ABA-JA and ABA-SA hormonal crosstalk could also be important for ABA responses in the plant-herbivore interplay. With respect to stomatal aperture, SA and methyl-JA can promote stomatal closure while other jasmonates, particularly the bioactive JA-Ile, can induce stomatal opening (Munemasa et al., 2011), though the contribution of SA and JA signalling might be secondary to ABA signalling (Zamora et al., 2021).

Most reports on the interaction between plants and arthropod herbivores have focused on insects, but mites are also highly economically relevant agricultural pests (Vacante, 2016). Among them, the two-spotted spider mite, *Tetranychus urticae,* an acari of the Tetranychidae family, is one of the most polyphagous pests found worldwide, feeding on more than 1,100 documented host plants, of which about 150 are crops (Migeon and Dorkeld, 2023). *T. urticae* pierces individual parenchymatic cells with stylets specialised for sucking their content. Infestation with *T. urticae* causes leaf chlorosis and severe infestation causes substantial crop losses (Bensoussan et al., 2016). In addition to its economic importance, *T. urticae* has a sequenced genome and a number of available tools and protocols (Grbić et al., 2011; Cazaux et al., 2014; Suzuki et al., 2017; Ojeda-Martinez et al., 2020). These characteristics and its ability to feed on the *A. thaliana* model plant led to *T. urticae* becoming a model herbivore for plant-pest interactions. However, potential positive or negative roles for ABA accumulation or signalling, including stomatal closure, in the context of plant defence against mite herbivores remain largely unexamined. How this key plant hormone that integrates a myriad of environmental cues impacts upon a generalist plant herbivore is relevant for a wider understanding of the plant-pest interplay.

Here, we have analysed the capacity of *T. urticae* to trigger stomata closure at the feeding site and on nearby undamaged tissues using *A. thaliana* as a host plant. We also studied stomatal behaviour and hormone crosstalk during mite infestation. We have demonstrated accumulation of ABA and characterised this accumulation at cellular resolution using the nuclear localised FRET biosensor for ABA nlsABACUS2-400n (Rowe et al., 2023). Mite feeding bioassays in mutants for ABA production and perception as well as in lines with altered stomata density have established the biological relevance of stomatal control, which are entry gates for mite stylets, for plant defences in the plant-mite interaction.

## Results

### Stomata responses to *T. urticae* infestation are driven by hormone content

To study time-resolved dynamics of defence and stomatal regulation upon mite infestation, stomatal aperture on the abaxial leaf surface was analysed in epidermal samples at four infestation time points (Fig 1A). Leaf epidermal impressions showed open stomata during the day with subtle reductions in aperture in wildtype Col-0 non-infested plants towards the end of the light period (Fig. 1A). Mite infestation induced a striking stomatal closure reaching the maximum closure at 24-30 h post-infestation (Fig. 1A, B). To also explore if natural stomatal closure in darkness could alter mite feeding habits and resultant leaf cell death, plants were grown in regular conditions, infested, incubated under constant dark or light environment for 16 h, and then cell damage was assayed. Results demonstrated that mites fed better under light conditions, when stomata were open, as plant cell death was increased (Fig. 1C).

**Figure 1.**
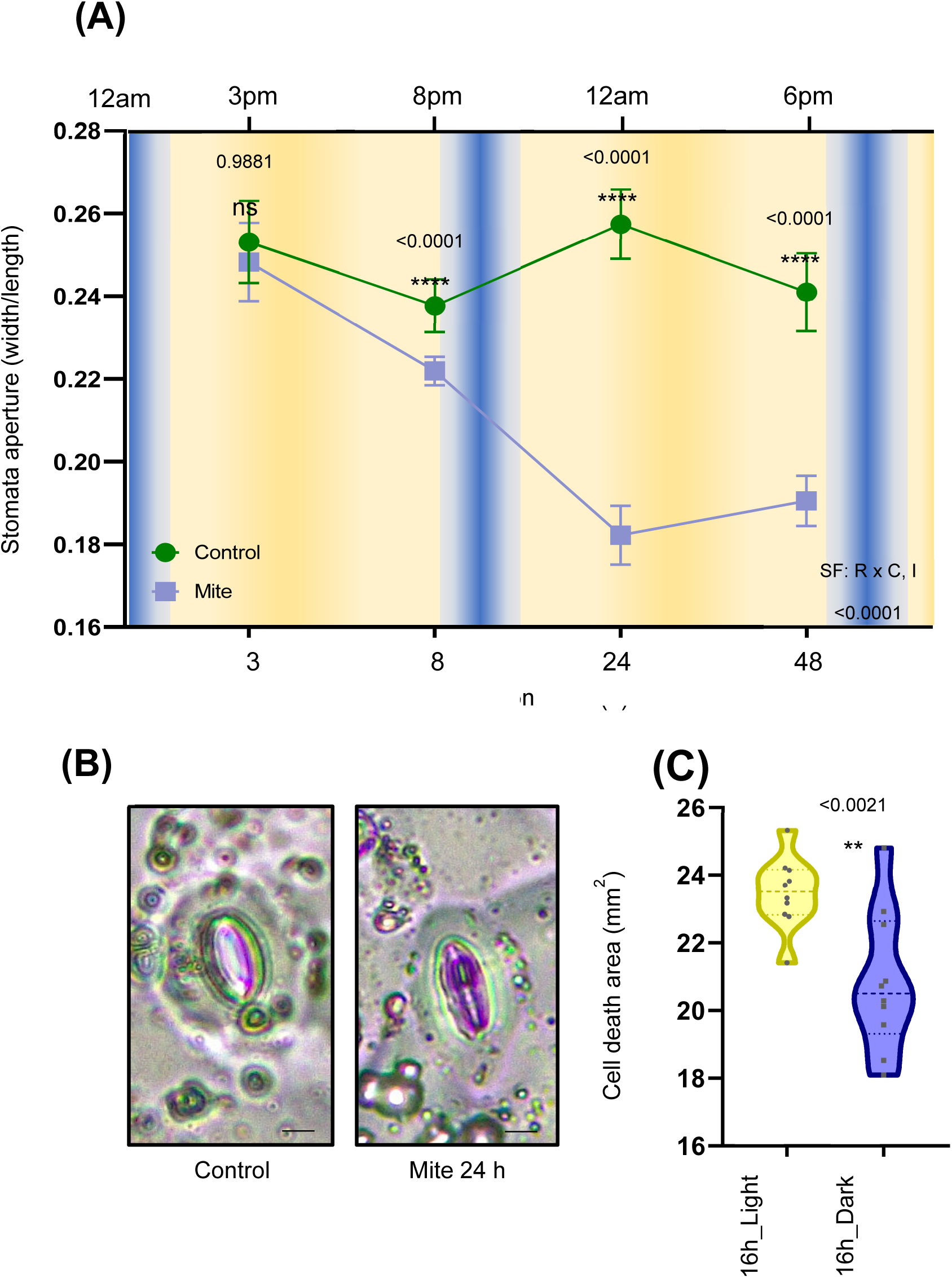
Effects of mite infestation on stomata aperture, and on the leaf cell death when plants are incubated under light or dark conditions. (A) Stomata aperture measured in Arabidopsis Col-0 after 3, 8, 24 and 30 h of mite infestation. Results referred as width/length ratio. Significant factors (SF) indicate whether the two independent factors, R (infestation time) and C (mite treatment), and/or their interaction I (RxC) were statistically significant (Two-way ANOVA followed by Student-Newman-Keuls test, P<0.05). Asterisks and numbers indicate significant differences between light and dark conditions. Data are means ± SE. (B) Image of stomata closure 24 h after mite infestation. Bars = 8µm (C) Cell death, measured by trypan blue staining after 16 h of mite infestation under dark and light conditions, is expressed in mm2. t-Student test was accomplished to assess differences due to light treatments (P<0.05) marked with one, two or three asterisks depending on significance. Data are means ± SE.

Since hormone accumulation in response to mites might be responsible for phenotypic changes in stomatal behaviour, SA, JA, JA-Ile and ABA were quantified in mite infested Arabidopsis Col-0 leaves at four post-infestation time points. The accumulation of all four compounds was induced by mites, but with distinct temporal profiles. SA content was significantly increased from the earliest infestation time while JA, JA-Ile and ABA required longer times to be differentially accumulated in comparison to non-infested plants (Fig. S1A-D). The highest hormone levels were detected at 24 h of infestation, except for the JA-Ile, which reached the highest content at 30 h of infestation, in accordance with a reduction in its JA precursor at this time point. Interestingly, ABA levels were maintained at 30 h, while SA was reduced. To determine the impact of SA, JA and ABA on leaf stomatal aperture, these hormones were exogenously applied to Arabidopsis Col-0 plants. SA and ABA treatment triggered stomatal closure in treated leaves, as expected (effect size in Supplemental Table S1), though the effect of JA was negligible (Fig. S1E-G).

### ABA accumulation detected upon mite infestation

Since ABA accumulates in leaves during mite infestation concomitantly with stomatal closure, we sought to identify in which cells ABA could be functionally relevant for Arabidopsis defence using the high-resolution nlsABACUS2-400n FRET-based biosensor for ABA (Rowe et al., 2023). In leaves of Arabidopsis plants expressing the biosensor, nuclei of epidermal and internal cell types showed sufficient biosensor expression for segmentation (Fig. 2A, upper images). In all detected cell types, nuclear ABACUS2-400n fluorescence emission ratios were higher in infested plants than in controls (Fig. 2B, bottom images, 2A), indicating higher ABA concentrations. Mean emission ratio for all cells was significantly higher for mite infested leaf images (Fig. 2B). To better understand the cellular distribution of ABA accumulation, we used a nuclear morphology classifier based on the “FRETENATOR” tool to analyse cell type emission ratios. Nuclei were classified into five cell type groups: stomata, pavement, spongy mesophyll, bundle sheath, and vascular bundle cells. With infestation, the signal increased in all cell type groups but stomatal and vascular bundle cell groups showed the highest emission ratios (Fig. 2C, D).

**Figure 2.**
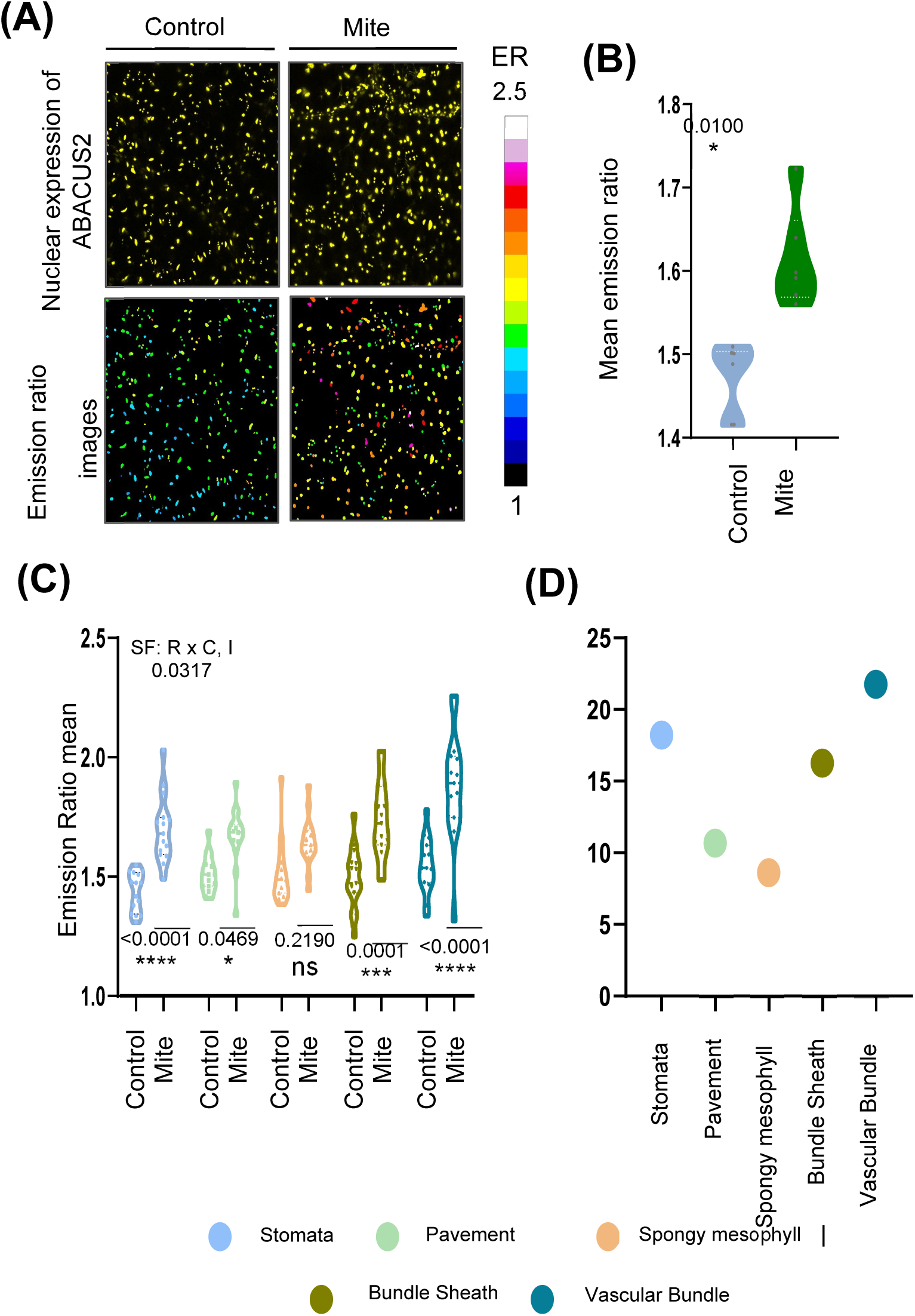
ABA accumulation at subcellular resolution level in leaf tissues of nuclear ABA-biosensor plants nls-ABACUS2-400n, after mite infestation. (A) Projections of analysed images. Upper images correspond to nuclei expressing the biosensor and bottom images to FRET-processed results. (B) Maximum Z-projections of the data quantified in (A). Upper images correspond to nuclei expressing the biosensor and bottom images to Emission ratios. t-Student test was accomplished to assess differences due to control and mite conditions (P<0.05) marked with one, two or three asterisks depending on significance. Data are means ± SE. (C) Emission ratios of nlsABACUS2-400n biosensor in cells of Vascular Bundle, Bundle Sheath, Spongy Mesophyll, Pavement and Stomata type cells, after 24 h of mite infestation. Significant factors (SF) indicate whether the two independent factors, R (mite treatment) and C (nucleus type), and/or their interaction I (RxC) were statistically significant (Two-way ANOVA followed by Student-Newman-Keuls test, P<0.05). Data are means ± SE. (D) Increased levels of emission ratio in the different five cell types after infestation respect to non-infested ones.

Because *T. urticae* cause cell death by inserting their stylets either in between epidermal pavement cells or through stomatal pores to suck mesophyll cell contents (Supplemental Fig. S2A; Bensoussan *et al*., 2016), it was important to get information about the viability status of leaf cells after the infestation. The ABA biosensor allowed us to detect cell viability since nuclei detected by biosensor fluorescence reflect viable cells. The comparison between infested and non-infested plants showed no significant changes in the total number of quantified nuclei (Supplemental Fig. S2B, Supplemental Fig. S7A). However, when this comparison was independently done in each of the five cell type groups, the mesophyll cell type presented lower number of nuclei in the infested plants than in non-infested (Supplemental Fig. S2C-G). Taken together, these results corroborated that spongy mesophyll cells were the target tissue where mites suck nutrients upon their feeding and indicated that nlsABACUS2-400n biosensor was able to detect local ABA increases in response to mites in leaf cell types directly involved in feeding as well as in distal cells and cell types not directly involved in feeding, i.e. bundle sheath and vascular cells. Interestingly, vascular cells have been proposed to be the major sites of foliar ABA biosynthesis (Endo et al., 2008), and thus our data provide support for a model in which mite feeding triggers ABA synthesis in the vasculature and this results in the translocation of ABA across the leaf cell types.

### ABA is involved in plant defence against mites

To study how changes in endogenous ABA levels affect mite infestation, Arabidopsis mutants in ABA biosynthesis and catabolism, *aba2-1* and *cyp707a1cyp707a3,* respectively, which result in lower and higher ABA concentrations (González-Guzmán et al., 2002; Okamoto et al., 2009a) were selected to perform mite infestation. *aba2-1* lines showed significantly greater damage than control plants upon mite feeding for 4 days while the damaged area in *cyp707a1cyp707a3* line was lower than in Col-0 plants (Fig. S3A). Likewise, fecundity rates, measured as cumulative number of eggs after 36 h, were higher when mites fed on the mutant *aba2-1* mutant line than when they fed on Col-0 plants, and were significantly reduced in the *cyp707a1cyp707a3* line (Fig. S3B). These data indicated that ABA promotes plant defences against mite damage and antagonises mite oviposition. Additionally, two Arabidopsis ABA insensitive mutants, the *112458* ABA receptor sextuple mutant and *ost1-3* SnRK2 kinase mutant, were also examined to determine if the canonical ABA signalling pathway participated in ABA-dependent mite responses. The OST1 gene is preferentially expressed in guard cells and the vasculature (Mustilli et al., 2002), cell types that exhibited the most pronounced increase in ABA levels (Figure 2). While mite females laid a similar number of eggs in mutants as in control plants, both mutant lines showed more damage than Col-0 plants after infestation (Fig. S3 C, D). Overall, these results revealed that ABA signalling, including in the vasculature or stomata, is important for mite defence.

Because all cells respond with increased ABA, we sought to determine whether ABA accumulation had further impact on SA and JA, considered as the core hormones involved in plant defences. We examined the expression of *PR1* and *MYC2*, marker genes of SA and JA signalling, respectively, in the five Arabidopsis genotypes, at two infestation times. The expression of the *PR1* gene was induced at 8 h post-infestation and decreased 24 h after mite feeding in all studied genotypes. However, *PR1* levels were lower in *aba2-1* line and higher in *cyp707a1cyp707a3* line than in Col-0 plants at 8 h time point and displayed the opposite expression pattern at 24 h (Supplemental Fig. S4A). No differences in *PR1* behaviour were found between *112458*, *ost1-3* mutants and wild-type (WT) plants (Supplemental Fig. S4B). Together these results indicate that ABA could crosstalk with SA signalling positively early in mite infection and negatively later in infection. In contrast to *PR1*, which peaked at 8 h, *MYC2* presented the maximum induction at 24 h after infestation (Supplemental Fig. S4 C, D). In the five tested genotypes, only the *aba2-1* mutant line showed altered MYC2 expression, i.e. elevated at 24 h (Supplemental Fig. S4 C, D). The observation that the reduced ABA levels in the *aba2-1* mutant increased expression of both *PR1* and *MYC* genes at 24 h could be an indication of hormone crosstalk but could also be an indirect result of increased mite infestation in *aba2-1*. Certainly, the overaccumulation of *MYC2* transcript suggests that a positive crosstalk of ABA with JA signalling is unlikely to explain the role of ABA in mite resistance.

One physiological response associated with ABA is the production of Reactive Oxygen Species (ROS), in particular H_2_O_2_, a signalling molecule that regulates plant stress response genes (Li et al., 2022). To further confirm such an association, the H_2_O_2_ content was determined in detached leaves of the four Arabidopsis mutants and Col-0 plants after 24 h of mite infestation (Supplemental Fig. S5). The measurement of H_2_O_2_ concentration is represented as relative DAB staining units (Supplemental Fig. S5A) and demonstrated that *aba2-1* and *cyp707a1cyp707a3* infested lines accumulated higher and lower levels of H_2_O_2_, respectively, than Col-0 plants. In contrast, number of DAB deposits in the *ost1-3* mutant was only slightly reduced and was not altered in *112458* after mite feeding (Supplemental Fig. S5). As with JA signalling, a negative association between ABA levels and reactive oxygen species (ROS) production suggests ROS accumulation is unlikely to explain the role of ABA in resistance to mite infestation.

### The effects of exogenous ABA application on mite infestation

As demonstrated above, exogenous ABA application induces stomata closure (Fig. S1G). To elucidate if ABA treatment affected plant defences, we pre-treated nlsABACUS2-400n plants by spraying ABA on the leaves 3 h before mite infestation and analysed cellular ABA levels and leaf responses to mites. As expected, ABA pre-treated and mite-infested plants displayed higher nlsABACUS2-400n biosensor emission ratios, indicating higher ABA levels, and the highest emission ratios were observed in those plants that were first pre-treated and then infested (Fig. 3A). Both ABA pre-treatment and mite infestation triggered stomata closure, with no synergistic or additive effects observed when the treatments were combined (Fig. 3B). The exogenous application of ABA before mite infestation helped plant defences since they showed less damage than non-pre-treated plants, and provided positive effects on plant growth, as shown in measurements of the rosette area (Fig. 3C, D,E). Thus, ABA pre-treatment enhanced mite defences and favoured plant growth during infestation.

**Figure 3.**
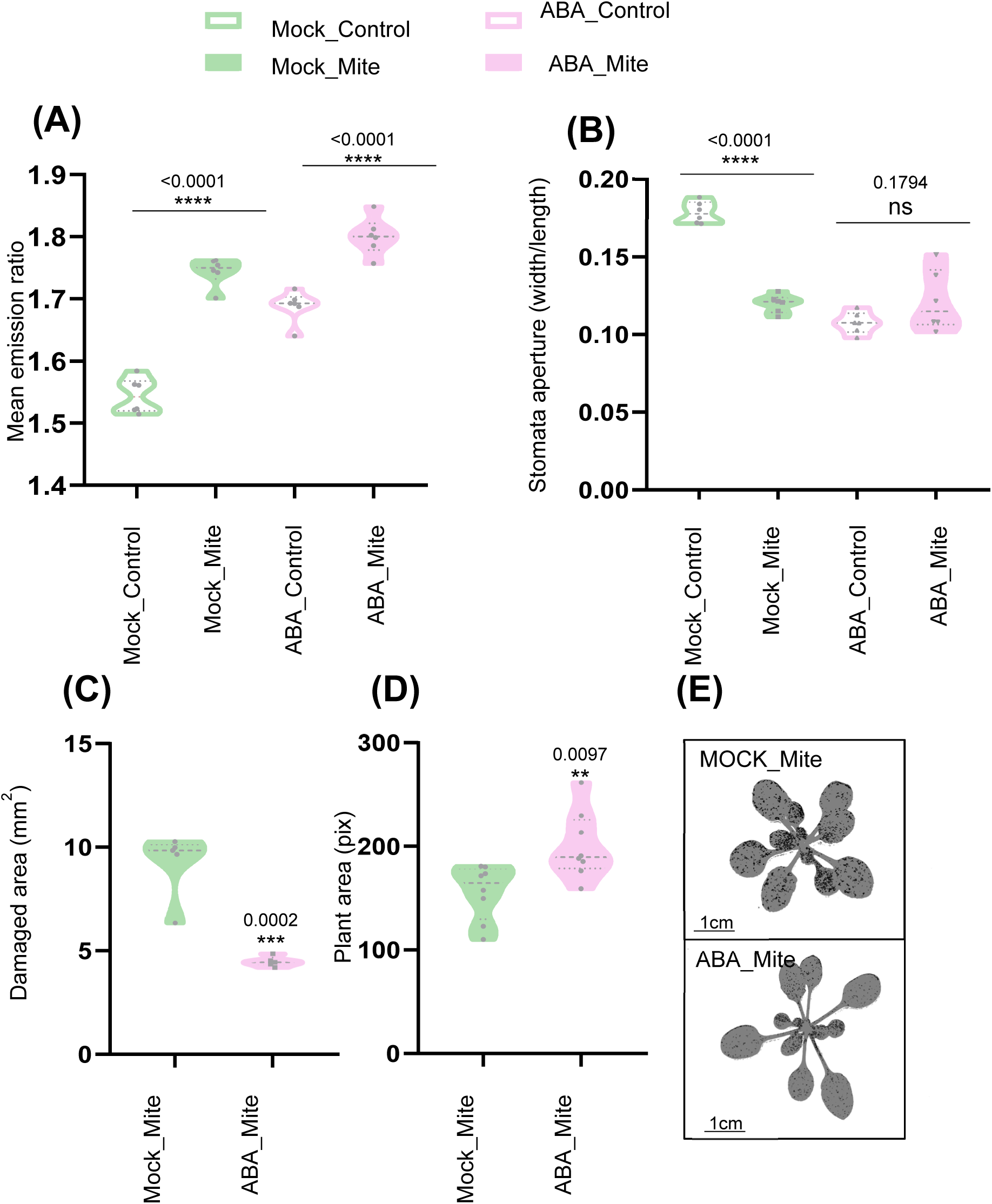
Effect of ABA pre-treatment of nls-ABACUS2-400n plants before mite infestation. Total emission ratio of biosensor signal (A), stomata aperture are referred to width/length the width/length ratio (B), damaged area expressed in mm^2^ (C), plant growth quantified in pixels (D) and infested Arabidopsis plant images ABA pre-treated and non-pre-treated (E), of nls-ABACUS2-400n leaves either ABA-pre-treated for 3 h or non-pretreated, and then mite infested for 24h or in non-infested. t-Student test was accomplished in all the panel. (P<0.05) marked with one, two or three asterisks depending on significance. Data are means ± SE.

### Stomatal aperture determines mite infestation outcome

As ABA accumulation and stomata closure are induced in response to mites, we questioned whether increased stomata aperture could provide any advantage for mite infestation. To clarify this point, Col-0 plants were pre-treated with FC, a compound that promotes stomata opening (Hunt et al., 2010), with ABA to close stomata, or with both. After treatments, stomatal aperture and plant damage were measured, and results were compared with mock-treated and non-infested plants. Plants pre-treated either with ABA or FC presented closed and opened stomata, respectively, and the combination of both produced a stomatal aperture comparable to mock plants (Fig. 4A, B). An experiment using different FC+ABA ratios demonstrated that very low concentrations of FC (0.5 µM) were sufficient to keep stomata open (Supplemental Fig. S6). Upon mite infestation, stomata remained closed in ABA treated plants, remained open in FC treated plants, and were actively closed similarly to mock when ABA and FC were previously combined (Fig. 4B). Plant damage results showed that more open stomata in the FC treatment correlated with higher leaf damaged area, indicating that open stomata facilitated mite feeding (Fig. 4C). As leaf damage increased in plants subjected to combined ABA and FC treatments compared with the ABA treatment alone, the control over stomatal aperture function of ABA is likely key for resistance to mite infestation.

**Figure 4.**
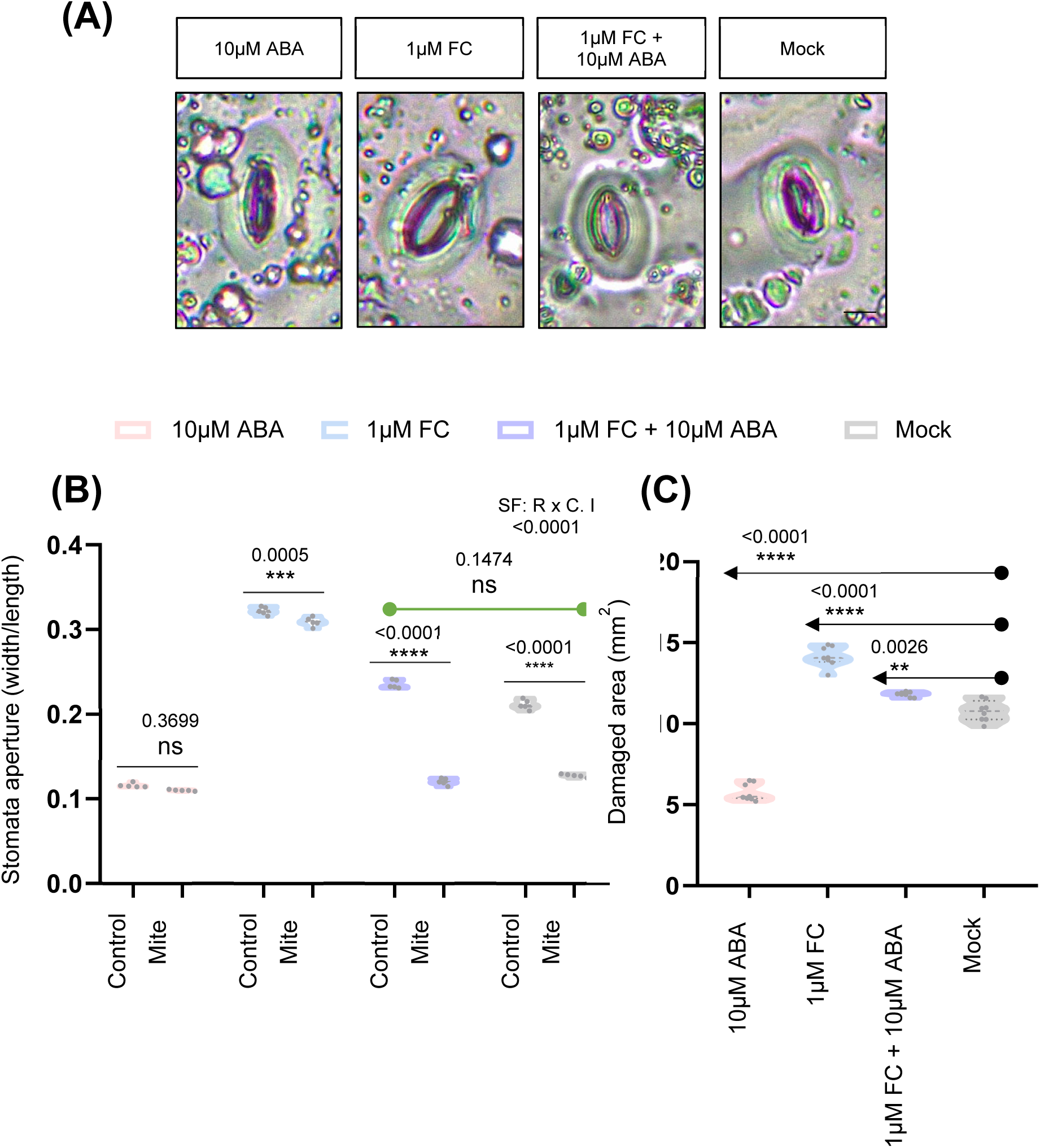
Effects of ABA (10 µM) and/or fussicoccin (1 µM) pre-treatments on stomata aperture and on Arabidopsis plant responses to mite infestation. (A) Images of stomata behaviour in ABA and/or fusicoccin (FC) pre-treated plants. Bars = 8µm (B) Stomata aperture in ABA and/or fusicoccin (FC) pre-treated plants after 24 h of mite infestation (C) Significant factors (SF) indicate whether the two independent factors, R (mite treatment) and C (Pre-treatment), and/or their interaction I (RxC) were statistically significant (Two-way ANOVA followed by Student-Newman-Keuls test, P<0.05). Plant damage of ABA and/or fusicoccin (FC) pre-treated plants after 4 d of mite infestation. Asterisks and numbers indicate significant differences compare to Mock treatment. Data are means ± SE.

### Effect of stomatal density on mite infestation

Given the significance of stomatal closure as a defence mechanism against mite infestation, we investigated the potential involvement of stomatal density in the leaf as a contributing factor using Arabidopsis lines with higher and lower number of stomata than Col-0 plants. The selected plants were mutants in epidermal patterning factors (EPF), a family of secreted peptides that inhibit stomatal development (Hepworth et al., 2015). In particular, we selected the *epf1epf2* mutant line and EPF2OE, an EPF2 over-expressing line. First, stomatal density was measured prior to mite infestation (Fig. 5A), and as expected, the *epf1epf2* mutant exhibited an elevated stomatal count, while the EFF2OE line displayed a reduced number of stomata. Then, we evaluated stomatal aperture in the three Arabidopsis genotypes with and without mite infestation and found that stomatal aperture in EPF2OE plants was higher than in *epf1epf2* or Col-0 plants (Fig. 5B). Interestingly, infested *epf1epf2* mutant exhibited somewhat lessened stomatal closure than Col-0, though infestation closed stomata in all genotypes (Fig. 5B). As stomata allow gas exchange between leaf mesophyll cells and atmosphere, contributing to leaf cooling (Chowdhury et al., 2021), we also analysed whether the leaf temperature was dependent on the stomata number and if temperature variations were produced during mite infestation. Uninfested temperature of *epf1epf2* leaves was 2.2 °C less than Col-0 and 3.1 °C less than EPF2OE leaves (Supplemental Table S4, Fig. 5C), consistent with stomatal density measurements. After mite infestation, an expected increase in leaf temperature was detected in Col-0 (+2.2 °C) and to a lesser extent in EPF2OE (+1 °C) plants that have fewer stomata. Mite infestation augmented leaf temperature in *epf1epf2* (+3.0 °C), the mutant line with more stomata, although infested leaves remained cooler than in infested Col-0 and EPF2OE (Supplemental Table S2, Fig. 5C). These temperature data corroborated the interrelationship between stomatal aperture and density in the plant response to mite infestation.

**Figure 5.**
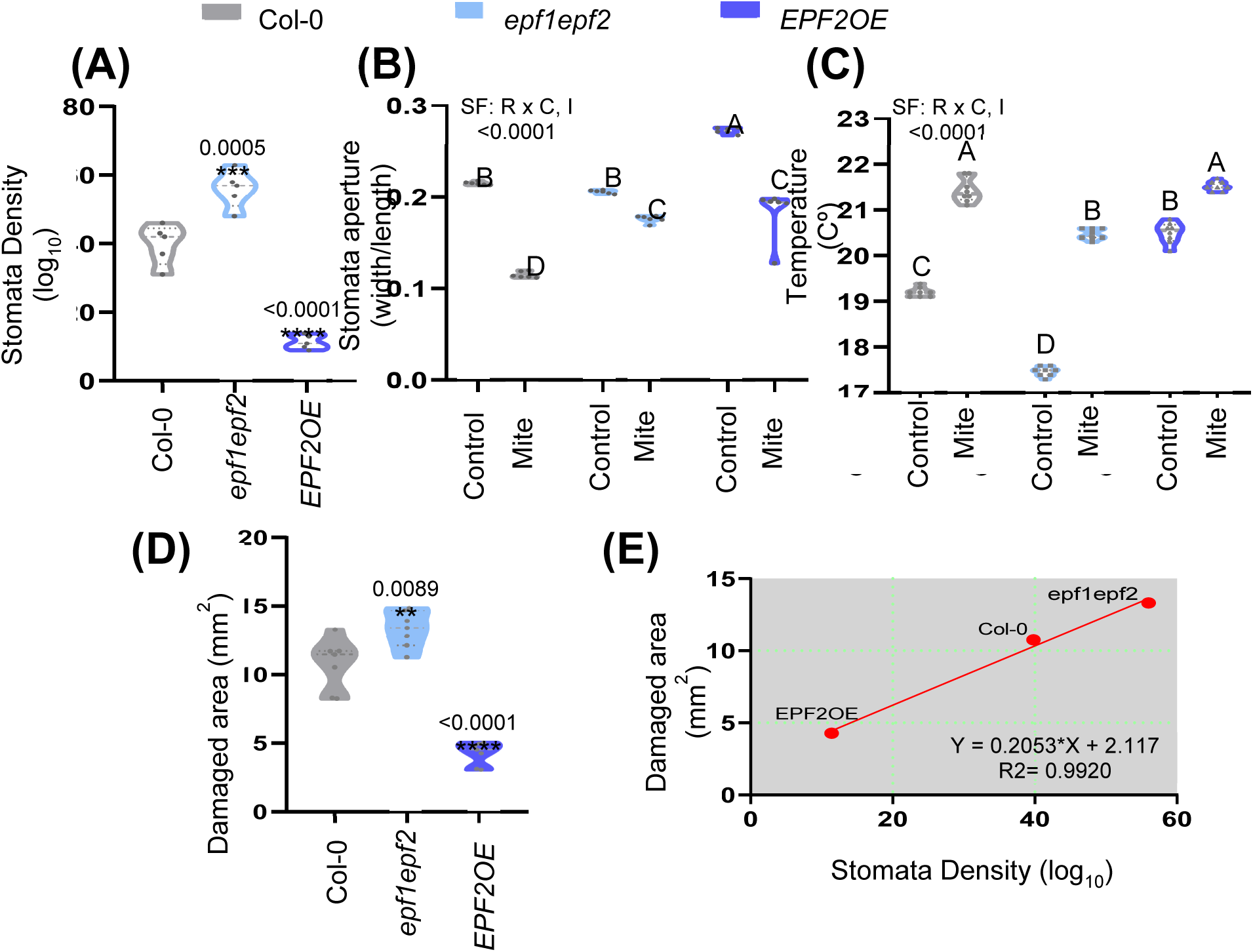
Stomata density in *epf1epf2* mutant, *EPF2OE* over-expressing line and Col-0 plants, and determination of damage, stomata aperture and temperature in the three genotypes after mite infestation. (A) Stomata density is expressed as Log_10_. (P<0.05) marked with one, two or three asterisks depending on significance (B) Stomata pore aperture are referred to the width/length ratio. (C) Temperature in °C. Significant factors (SF) indicate whether the two independent factors, R (mite treatment) and C (genotype), and/or their interaction I (RxC) were statistically significant (Two-way ANOVA followed by Student-Newman-Keuls test, P<0.05). Data are means ± SE. (D) Foliar damage, expressed in mm^2^, was quantified 4 d after mite infestation. t-Student test was accomplished. (E) Correlation between stomata density and damaged area for the three Arabidopsis genotypes. Data are means ± SE.

Finally, the damage produced by mites (Fig. 5D) was quantified in these Arabidopsis plants. The *epf1epf2* line displayed more damage than Col-0 plants, while in the EPF2OE line the mite damage was significantly reduced. The close correlation between stomatal density and damage (R squared = 0.9920; Fig. 5E) indicate that the density of stomata in leaves is an essential feature that determines the success of mite infestation.

## Discussion

Stomata are specialized microscopic gates at the epidermal plant surface with the ability to be opened or closed in response to environmental and endogenous signals. Stomatal movement depends largely on the content of ABA, considered as the key hormone that closes stomata, although other hormones also participate in governing stomata responses (Wei et al., 2021). Under a subset of biotic stresses, ABA levels and stomata behaviour play an essential plant defence role by hindering adversary access to the leaf interior (Lim et al., 2015; Melotto et al., 2017). In the plant-mite context, stomata are target sites where mites insert their stylets to gain access to nutrient rich mesophyll cells. Once the plant perceives the mites, a signal transduction cascade is triggered that activates the synthesis of JA and SA, besides other chemical responses, to generate defences (Zhurov et al., 2014; Santamaría et al., 2019). Our results confirmed that mite infestation induced JA and SA, demonstrated the timeline of this response, and also revealed a mite induced increase of ABA. Other studies have linked stomata movement with response to light and circadian-clock pathways (Tallman, 2004) and we observed that *T. urticae* caused less cell damage by feeding under dark conditions when stomata are closed. We also observed stomatal closure concomitant with ABA accumulation in response to herbivory during the day. Additionally, the temporal pattern of the phenotypic changes in stomatal behaviour in response to mite infestation during the day matched with the ABA accumulation more than the JA or SA content. Together, these data pointed to ABA accumulation and stomatal closure as a plant response to herbivory acting to limit leaf cell damage associated with mite infestation.

Stomatal closure has been previously described as a plant response to other phytophagous species (Sances et al., 1979; Pincebourde and Casas, 2006; Schmidt et al., 2009), though whether closure constituted a defence mechanism or contributed to the infestation remained unclear. DNA microarray experiments of Arabidopsis infiltrated with *Myzus persicae* saliva allowed the identification of a number of aphid saliva up-regulated genes that also responded to ABA treatment (Hillwig et al., 2016). Aphid infestation induced ABA production in Arabidopsis, but aphids showed preference for and performed better on wild-type plants than on ABA deficient mutants, suggesting that ABA might not be a plant defence response. These authors also demonstrated that ABA increased the synthesis of some glucosinolates with defensive properties. We demonstrated a positive role for ABA in the plant defence to the mite *T. urticae* since mutant lines in ABA biosynthesis and catabolism showed higher and lower leaf damage, respectively, than Col-0 infested plants. These data directly corresponded to the higher and lower number of eggs accumulated in these mutants. Thus, mite infestation results in ABA accumulation as a plant defence response that closes stomata, reduces mite feeding ability, lessens leaf damage and lowers mite fecundity rates.

Vos *et al*., (2013) also reported that ABA is a crucial regulator of induced resistance against *Pieris rapae* by activating primed JA-regulated defence responses in Arabidopsis. Additionally, ABA has also been associated with redox homeostasis (Li et al., 2022), in particular with ROS accumulation, which is also essential in the plant defence against mites (Santamaría et al., 2017; Arnaiz et al., 2021). Our results showed increased levels of JA signalling and H_2_O_2_ production in ABA deficient mutants after mite infestation where leaf damage was increased. Based on the current observations, it can be concluded that ABA signalling, independent of positive crosstalk with ROS and JA signalling, plays a critical role as a defence determinant against mite infestation.

In order to probe the regulation and function of ABA accumulation, we used nlsABACUS2, a FRET-based biosensor for ABA, as a tool to quantify *in vivo* ABA at high spatio-temporal resolution in response to mite infestation. Elevated levels of ABA were broadly detected in the nuclei of a range of leaf cell types, with slightly stronger responses in stomata and vascular tissues. These ABA dynamics are consistent with an ABA regulation model in which the leaf vasculature is the key site of biosynthesis triggered by mite infestation with ABA subsequently transported to guard cells for stomatal closure, similar to that proposed for ABA during water stress by Kuromori *et al*., (2018). However, we also show induction of ABA in all leaf cells involved in feeding by mites, including pavement cells and the mesophyll cells on which they feed.

Given the pronounced stomatal closure observed during infestation, an additional significant aspect of this study focused on clarifying the impact of stomatal status versus ABA signalling more generally. This was achieved by administering exogenous ABA or FC, to induce or repress stomatal closure. When pre-treated with ABA and subsequently infested with mites, plants were protected without sacrificing their growth. Pre-treatment with FC increased leaf damage, consistent with a protective role for stomatal closure. In an ABA + FC treatment, mite damage correlated with stomatal aperture rather than ABA levels, providing further support for the role of stomata in ABA mediated mite defence. The closure of stomata likely hindered mite feeding through the natural openings on the leaf, compelling them to utilize their stylets for penetrating closed stomata or between the epidermal pavement cells. This mode of stylet penetration could trigger earlier damage and possibly defence signalling and likely reduces feeding success since mites try to avoid epidermal cell damage (Bensoussan et al., 2016) and we observed ABA to reduce overall leaf damage and defence signalling at 24 h infestation.

We also investigated the importance of leaf stomatal density upon infestation based on the hypothesis that higher number of stomata could facilitate mite feeding. In mutants with more and less stomata, we observed lower and higher leaf temperature that increased with mite feeding, indicating that mite induced stomatal closure remained functional in leaves with altered stomatal density. Indeed, stomatal density was tightly correlated with mite induced leaf damage, suggesting that the number of stomata, in conjunction to their aperture, is a highly relevant trait for plant resistance to mite infestation.

This study provides valuable insights into the important role of stomata and its regulation mediated by ABA in the Arabidopsis defence against *T. urticae*. We demonstrate that ABA, possibly of vascular origin, accumulates in the guard cells of the stomata as a result of mite feeding and triggers stomatal closure as a structural defence response. We also show stomatal aperture and abundance play a crucial role in determining the success of infestation. As both traits are regulated by ABA, the levels of this phytohormone, which also integrate myriad other biotic and abiotic stress cues, is an important determinant of mite infestation that could be considered in breeding programs for pest control.

## Material and methods

### Plant material and growth conditions

*Arabidopsis thaliana* ecotype Col-0, from Nottingham Arabidopsis Seed Collection (NASC; http://arabidopsis.info/BasicForm/), was used in all experiments as wild-type (WT). Plants expressing nuclear localized ABA FRET biosensors (nlsABACUS2-400n (Rowe et al., 2023) were used to quantify cellular ABA accumulation during mite feeding assays. *Aba2-1* (González-Guzmán et al., 2002), *cyp707a1 cyp707a3* (Okamoto et al., 2009b), *pyr1pyl1pyl2pyl4pyl5pyl8 (*hereafter *112458)* (Gonzalez-Guzman et al., 2012), *ost1-3* (SALK_008068) and *epf1epf2* mutant lines and EPF2OE (Hara et al., 2009; Hunt et al., 2010) genotypes were used for ABA, stomata and mite feeding assays. All plants were grown in a mixture of peat moss and vermiculite (2:1). Sterilized seeds were stratified in the dark at 4°C for 5 days as in Santamaria *et al*. (2019). Plants were then grown in growth chambers (Sanyo MLR-351-H) under controlled conditions (23°C±1°C, >70% relative humidity and a 16 h:8 h light:dark photoperiod). The growth chamber includes a light control program to mimic sunrise and sunset conditions.

### Spider mite growth, maintenance and plant infestation

*Tetranychus urticae* London strain (Acari: Tetranychidae) population, provided by Dr. Miodrag Grbic (UWO, Canada), was reared on *Phaseolus vulgaris* (beans) and maintained in growth chambers (Sanyo MLR-351-H, Sanyo, Japan) at 25°C±1°C, >70% relative humidity and a 16 h:8 h (light:dark) photoperiod. Mite infestation was performed in 3-week-old Arabidopsis rosettes or in detached leaves collected from the corresponding Arabidopsis genotypes with 20 mites/plant or with 10 mites/leaf, adapting the infestation time according to the experiment. For leaf infestation assays, leaf number 5 or 6 were selected (Merchant & Pajerowska-Mukhtar, 2015), placed on a one-half-strength Murashige and Skoog (MS) (Duchefa Biochemie) medium plates, infested and covered but ventilated with a relative humidity of 60-80%.

### Hormone analysis

Frozen plant material (c.a. 50 mg) was used for the extractions. Plant samples were spiked with 25 µl of an internal standard mixture (containing ABA-*d_6_*, DHJA and C^13^-SA) to correct for analyze losses as in (De Ollas et al., 2021). Extraction was carried out in ultrapure water in a ball mill at room temperature using 2 mm glass beads. Homogenates were centrifuged at 10,000 rpm for 10 min at 4°C and supernatants recovered. The resulting solutions were partitioned twice against an equal volume of diethyl ether after adjusting pH to 3.0 with a 30% acetic acid solution. The combined organic layers were evaporated under vacuum in a centrifuge concentrator (Jouan, Sant Germaine Cedex, France) and the dry residues reconstituted in a 10% (w/v) aqueous methanol solution. Prior to injection, extracts were filtered through 0.20 µm PTFE syringe membrane filters and filtrates recovered in chromatography amber glass vials. Samples were analyzed by tandem LC/MS in an Acquity SDS UPLC system (Waters Corp., USA) coupled to a TQS triple quadrupole mass spectrometer (Micromass Ltd., UK) through an electrospray ionization source. Separations were carried out on a C18 column (Luna Omega Polar C18, 50×2.1 mm, 1.6 µm particle size, Phenomenex, USA) using a linear gradient of ultrapure acetonitrile and water, both supplemented with formic acid to a 0.1% (v/v) concentration, at a constant flow rate of 0.3 mL min^-1^. During analyses, column temperature was maintained at 40°C and samples at 10°C to slow degradation. Plant hormones were detected in negative electrospray mode following their specific precursor-to-product ion transitions and quantitated using an external calibration curve with standards of known amount.

### Stomata leaf impression and aperture quantification

Stomata leaf impressions were done using detached leaves, either treated and/or mite infested (after a previous mite removing). Leaves were pressed on fast-setting dental resin AquasilUltra+ (Dentsply Sirona) which was allowed to set, then leaf material was removed. Transparent nail polish was spread onto the resin pieces (Wang et al., 2006). After they had thoroughly dried, nail varnish impressions were peeled using Sellotape and affixed to a glass microscope slide. These slides were then imaged on a Leica DM1000 LED with an ICC50 W camera. Stomatal density and aperture were quantified with Fiji ImageJ software (Schindelin et al., 2012).

### Cell death quantification

Cell death quantification was performed by trypan blue staining after 16 h of infestation in light and dark conditions. Leaf discs were boiled in trypan blue solution followed by a clarification process with 2.5 g/ml of chloral hydrate (Sigma) solution Sanchez-Vallet *et al*. (2010). Discs were placed onto glass slides in 50% (v/v) glycerol and observed under an epifluorescence stereoscope using UV filters. Quantification was performed with Pixel quantification was performed using Adobe Photoshop (Luna et al., 2012).

### Exogenous plant pre-treatments

Exogenous treatments were done on the aerial part of the plant by spraying with 1 mM SA, 1 mM JA, 10 µM ABA or 1 µM Fusicoccin (FC) (all from Sigma) dissolved in absolute ethanol. After 3htreatment, mite infestation or stomata leaf impressions were performed.

### Confocal microscopy and images processing

An inverted SP8 confocal microscope (Leica) was used for biosensor imaging assays. All images were acquired as Z-stacks in 16-bit mode, with a 10X dry objective. Samples were mounted in ¼ MS, pH 5.7. Typical settings were as follows: sequential scanning was performed with excitation lasers and HYD detectors. 442 nm excitation 3-10% was used with– HYD1: 460-500 nm, 100 gain for a first acquisition to detect the donor T7edCerulean fluorescence (donor excitation, acceptor emission or DxAm). Second, 442 nm excitation 3-10% was used with HYD2 525-560 nm, 100 gain to acquire energy transfer fluorescence (donor excitation, donor emission or DxDm). Next, 514 nm excitation 5-10% was used with HYD2 525-560nm, 100 gain for a third acquisition to detect the edCitrineT7 acceptor protein fluorescence (acceptor excitation, acceptor emission or AxAm). Scan speed was set at 400, Line averaging: 2-4, Bidirectional X: on.

### FRET Cell and Tissue classification

The image processing for fluorescence emission ratio (DxAm/DxDm) quantification was carried out using the “FRETNATOR” tool reported by Rowe *et al*., 2022 and Rowe *et al*., (2023) (Fig. S5B). The AxAm images were used for segmentation of nuclei. The specific cell-type emission ratio classification was done by the FRETcelltype extension plugin. The extension was developed by using Apache Groovy and reads each series of acquisitions (Leica file format (.lif). FRETcell type uses the “FRETENATOR_Segment_and_ratio’’ output, specifically the “threshold image output” and “The label map” to classify the nuclei based on the ROI shape. Cell types were defined as Vascular Bundle, Bundle Sheath, Spongy Mesophyll, Pavement and Stomata cells. The plugin provides two tables (.csv) for each image. The “Result Table” and the “Summary Table”. The “Result Table” displays the number of pixels, the coordinates, and the emission ratio value for each nucleus Supplemental Fig. S7B provides information about the nuclei classification shape validation.

### Plant damage and mite performance assessment

Chlorotic damage in whole plants was quantified 4 days after mite infestation according to (Ojeda-Martinez et al., 2020), using nine biological replicates from independent rosettes for each genotype. Mite fecundity was performed in detaches entire leaves from different Arabidopsis genotypes, infested with ten synchronized females per leaf, and number of eggs were counted after 36 h, following (Santamaría et al., 2019).

### Nucleic acid analysis

Total RNA was extracted from Arabidopsis rosettes by the phenol/chloroform method, and precipitated with 8 M LiCl as described (Oñate-Sánchez and Vicente-Carbajosa, 2008). Complementary DNAs (cDNAs) were synthesized from 2 μg of RNA using the Revert AidTM H Minus First Strand cDNA Synthesis Kit (Fermentas). RTqPCR was performed using LightCycler® 480 SYBR® Green I Master (Roche), a SYBR Green Detection System (Roche) and the LightCycler®480 Software release 1.5.0 SP4 (Roche). mRNA quantification was expressed as relative expression levels (2-dCt) or as fold change (2-ddCt) (Livak and Schmittgen, 2001). Arabidopsis ubiquitin 21 was used as a housekeeping control gene and PR1 (Pathogenesis-Related protein-1) and *MYC2* (MYC2 transcription factor) as marker genes of SA and JA hormone signaling, respectively. Primer sequences are stated in Supplemental Table S3.

### Hydrogen peroxide determination

The accumulation of H_2_O_2_ was visualized in detached leaves after 24 h of mite infestation, using 3,3-diaminobenzidine tetrachloride hydrate (DAB, Sigma) as a substrate, according to (Martinez De Ilarduya et al., 2003). Discs were placed onto glass slides in 50% (v/v) glycerol and observed under a stereoscope. Pixel quantification was performed with Fiji ImageJ software (Schindelin et al., 2012).

### Thermal imaging determination

Thermal images were obtained using a FLIR® infrared camera (FLIR-T600) equipped with a 16° lens, 24 h after infestation. To avoid changes in leaf temperature due to environmental conditions, experiments were conducted inside a walk-in growth chamber set at 25°C, 60 ± 10% relative humidity, and light intensity of 105 μmol.m−2.s^−1^. The camera was vertically mounted at approximately 20 cm above the set up. Images were saved as 8-bit TIFF files and pictures were analysed using ImageJ (Fiji), in which all pictures were set at constant range of temperature based on pictures of Col-0 control conditions, emissivity of the samples was 0.925.

### Statistical analysis

Statistical analyses were done using GraphPad Prism v9.4.1. The normality and homoscedasticity of the data were previously analysed to apply the proper analysis. T-Student test was used for individual analysis. One-way ANOVA followed by Tuke’S multiple comparisons test was used to compare multiple data set. Two-way ANOVA was performed in the experiments that row data (R) and treatment with column data (C) were simultaneously analysed, and Tuke’s multiple comparisons test was used when the interaction (RxC) was significant. Correlation between damage and stomata density was analysed applying Pearson Product Moment correlation test. Statistical test applied are in Supplemental Table S4.

## Author contributions

AMJ, IRD and ID conceived the research. IRD, JR, ACL and VA performed the experimental research. ID and IRD wrote the first draft of the manuscript. All authors contributed to the final version of the manuscript.

## Acknowledgements

We thank Julie Gray and Eiji Nambara for providing ABA mutant seed. We also thank Marino Rodriguez-Exposito for his technical assistance.

## Code availability

FretCellType image analysis tool is available at: https://github.com/acayuelalopez/FretCellType

## Supporting Tables

**Table S1.** Effect size between SA. JA and ABA treatments in Arabidopsis plants while measuring stomata aperture.

|  | SA | JA | ABA |
| --- | --- | --- | --- |
| Mean of Mock | 0.2574 | 0.2577 | 0.2557 |
| Mean of Hormone | 0.1766 | 0.2513 | 0.1114 |
| Difference between means (Hormone - Mock)<br>± SEM | -0.08073 ±<br>0.003475 | -0.006482 ±<br>0.002826 | -0.1444 ±<br>0.002810 |
| 95% confidence interval | -0.08769 to -<br>0.07376 | -0.01215 to -<br>0.0008169 | -0.1500 to -<br>0.1387 |
| R squared (eta squared) | 0.909 | 0.0888 | 0.98 |

**Table S2.** Leaf temperatures in Arabidopsis mutants con different stomata density after mite infestation. Data are mean ± SE.

| Treatment | Mean (°C) |  |  |
| --- | --- | --- | --- |
|  | Col-0 | *epf1epf2* | *EPF2OE* |
| Control | 19.2 ± 0.038 | 17.4 ± 0.037 | 20.5 ± 0.080 |
| Mite | 21.4 ± 0.092 | 20.4 ± 0.041 | 21.5 ± 0.037 |

**Table S3.**
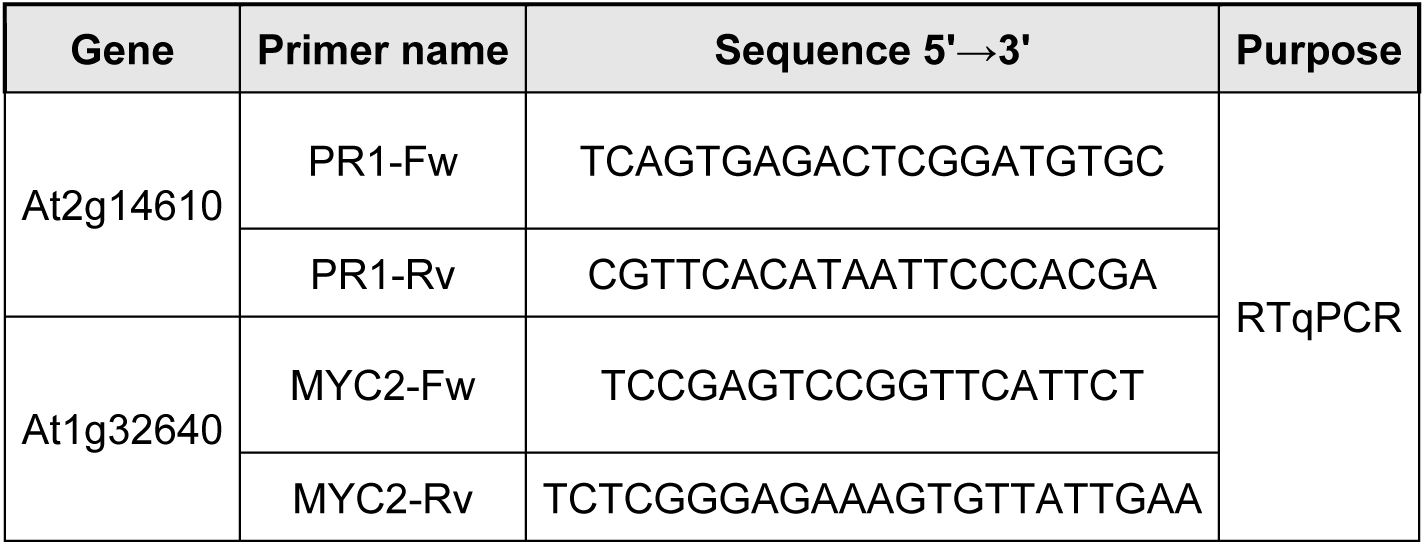
Oligonucleotide sequences. Primer sequences used for RTq-PCR assay.

**Table S4.**
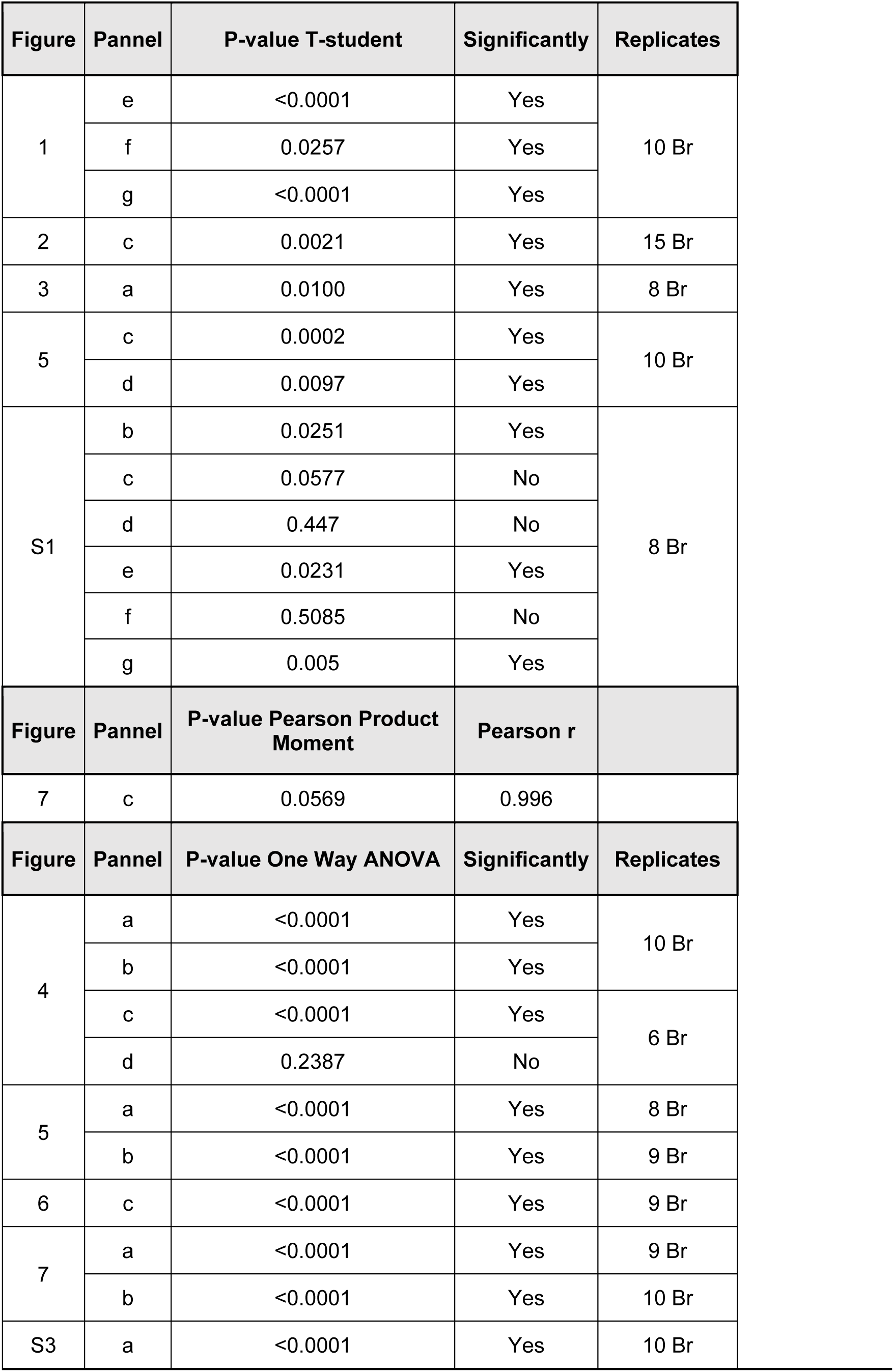
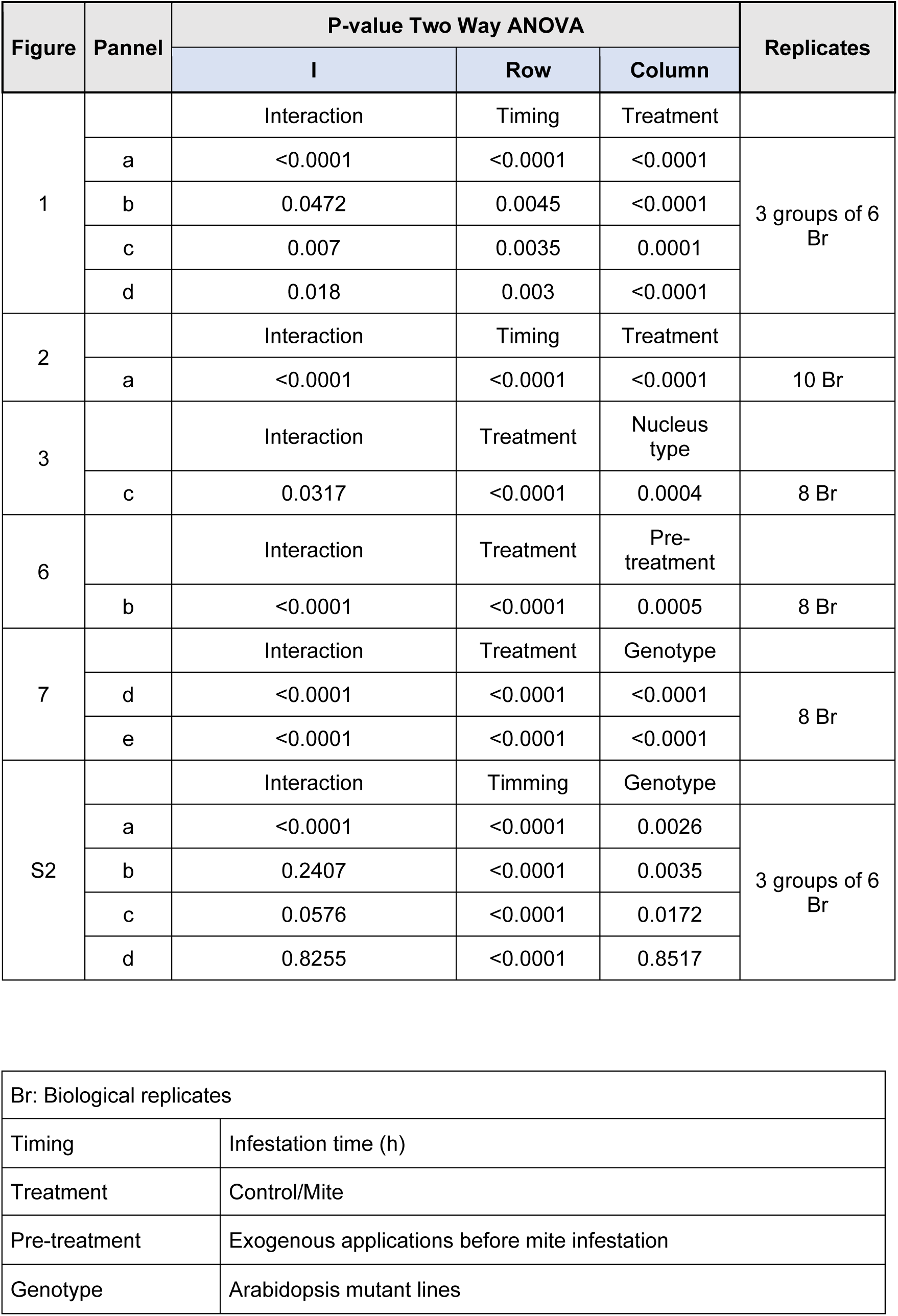
Statistical test applied indicating the corresponding figures.

