## Supplemental Figures for "Stomata, a vulnerability in the plant defence against phytophagous mites that ABA can overcome"

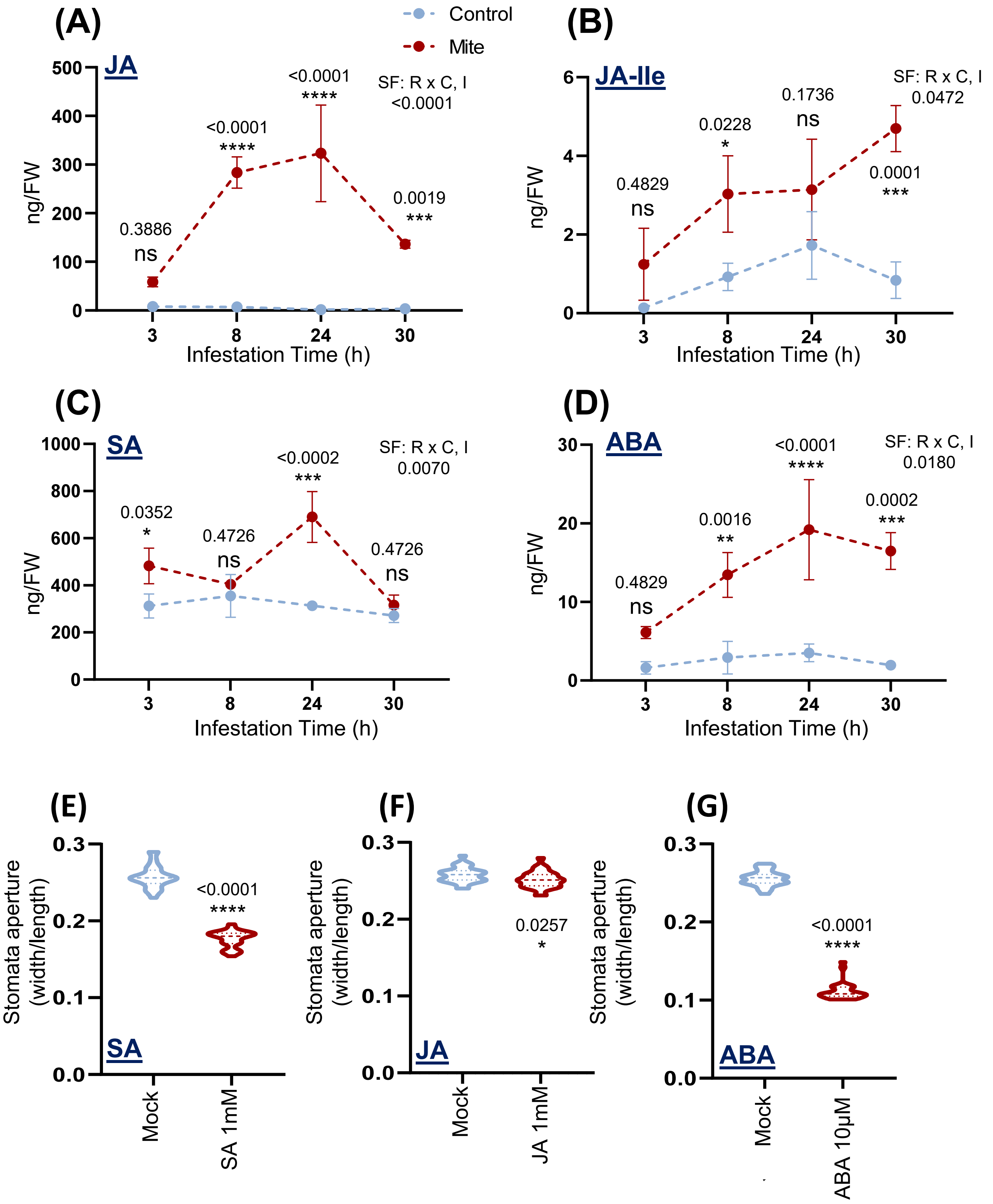


**Figure S1. Quantification of hormone content in Arabidopsis Col-0 upon mite infestation, and stomata aperture after exogenous hormonal treatments.** JA (A), JA-Ile (B), SA (C) and ABA (D) accumulation was quantified at 3, 8, 24 and 30 h after mite infestation. Values are expressed as ng of hormone per g of fresh weight (FW). Significant factors (SF) indicate whether the two independent factors, R (Infestation time) and C (mite treatment), and/or their interaction I (RxC) were statistically significant (Two-way ANOVA followed by Student-Newman-Keuls test, $P<0.05$). Asterisks and numbers indicate significant differences compare to control conditions. Data are means ± SE. Stomata aperture measured at 3 h after spraying on the aerial part of the plant with 1 mM SA (E), 1 mM JA (F) and 10 µM ABA (G). Results referred as width/length ratio. t-Student test was accomplished to assess differences due to hormone treatments ($P<0.05$) marked with one, two or three asterisks depending on significance. Data are means ± SE.

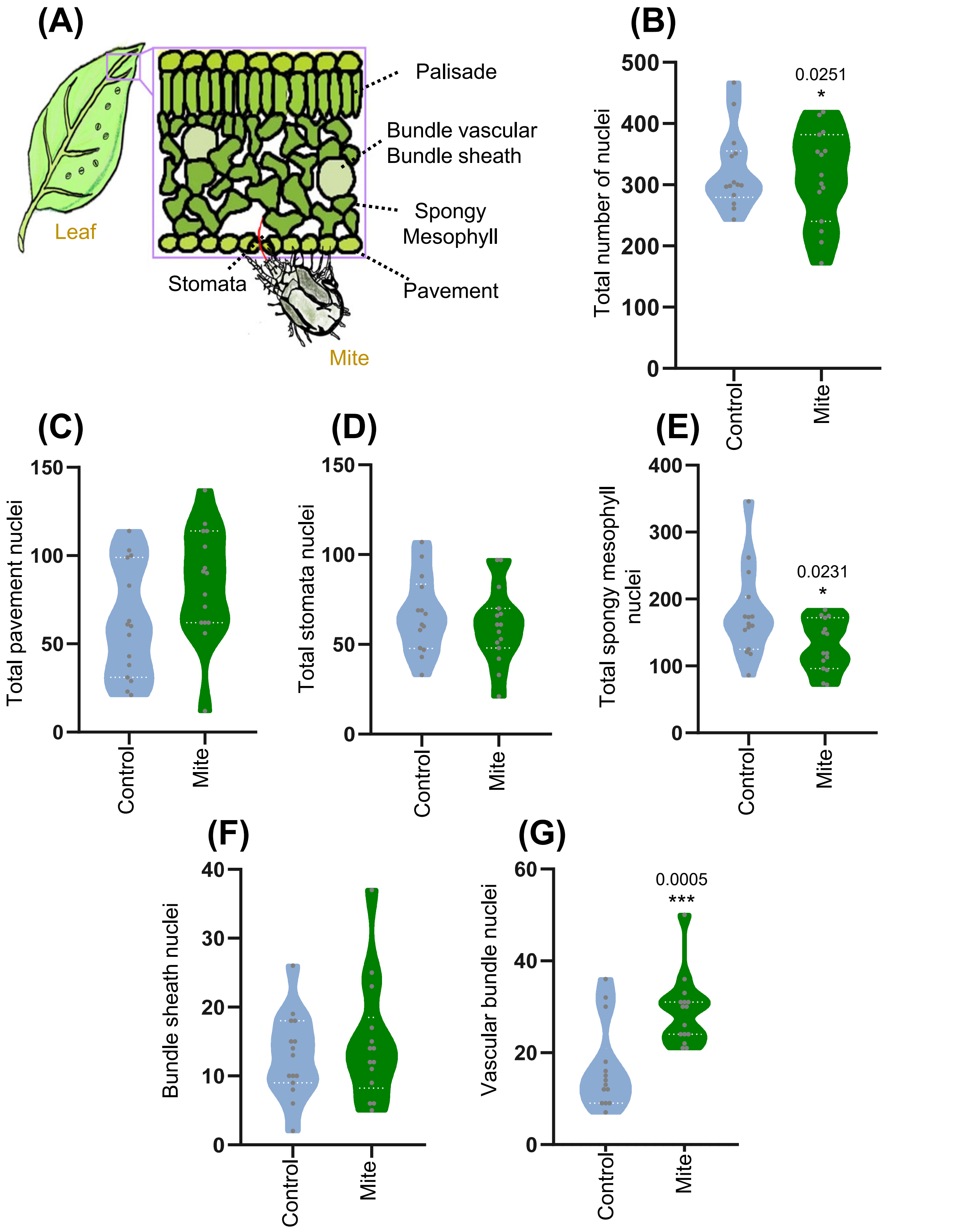


**Figure S2. Quantification of nuclei number in different cell type groups of nls-ABACUS2-400n plants after mite infestation.** (A) Scheme of the leaf tissues and a mite feeding through stomata. (B) Total number detected nuclei in infested and control leaves. (a) Total number of nuclei in different cell types: Pavement (C), Stomata (D), Spongy mesophyll (E), Vascular bundle (F) and Bundle sheath (G), after 24 h of mite infestation. Asterisks and numbers indicate significant differences between control and mite treatment.

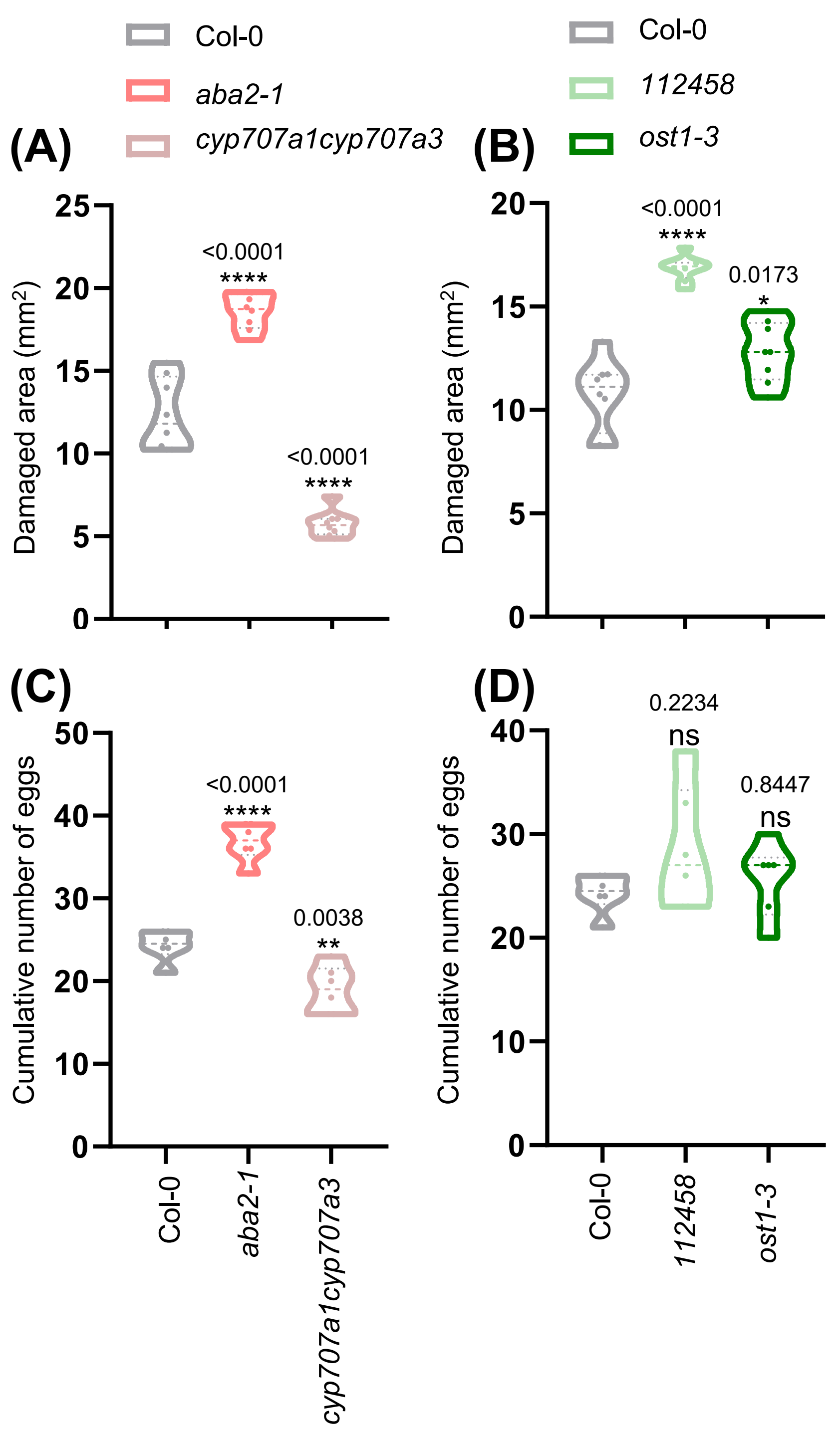


**Figure S3. Plant damage and mite fecundity after mite infestation of *aba2-1* and *cyp707a1cyp701a3* and *pyr1pyl-112458, ost1-3* Arabidopsis mutants and Col-0 plants.** Foliar damage quantified 4 d after mite infestation in *aba2-1*, *cyp707a1cyp707a3* and Col-0 plants (A), and in *112458, ost1-3* and Col-0 plants (B). Data, expressed in $mm^2$. Effects on *T. urticae* fecundity measured 36 h after the infestation with synchronized mite females on *aba2-1 cyp707a1cyp707a3* and Col-0 plants (C), and in *pyr1pyl-112458, ost1-3* and Col-0 plants (D). Asterisks and numbers indicate significant differences compare to Col-0 genotype. Data are means ± SE. One-way ANOVA followed by Newman-Keuls multiple comparisons test, P<0.05.

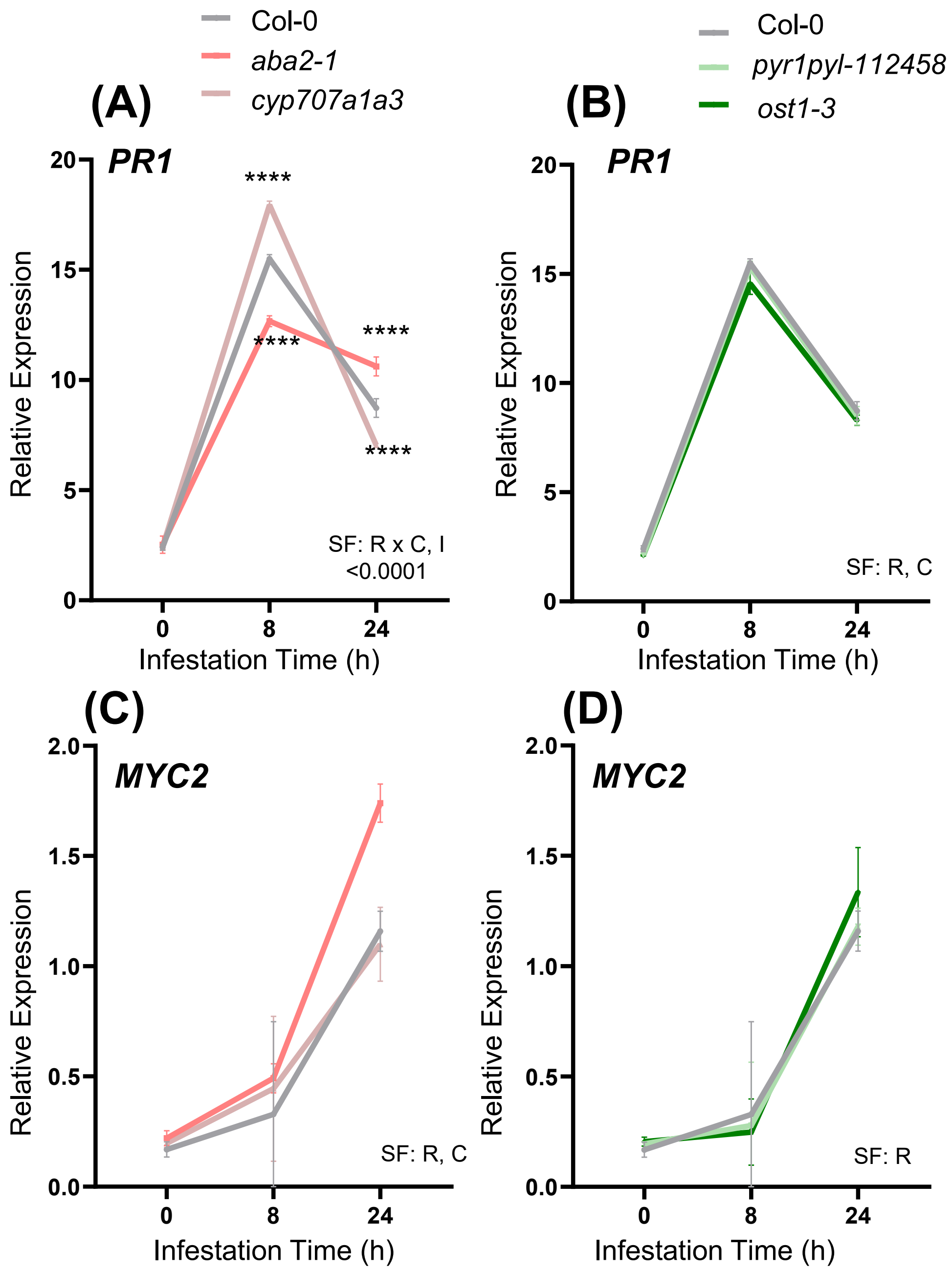


**Figure S4. Expression levels of *PR1* and *MYC2* genes in the five Arabidopsis genotypes.** *PR1* gene (A,B) and *MYC2* gene (C,D). Gene expression levels were determined in *aba2-1*, *cyp707a1cyp707a3,* and in *pyr1pyl-112458, ost1-3* and Col-0 plants. Values indicated as relative expression. Significant factors (SF) indicate whether the two independent factors, R (infestation time) and C (genotype), and/or their interaction I (RxC) were statistically significant (Two-way ANOVA followed by Student-Newman-Keuls test, P<0.05).

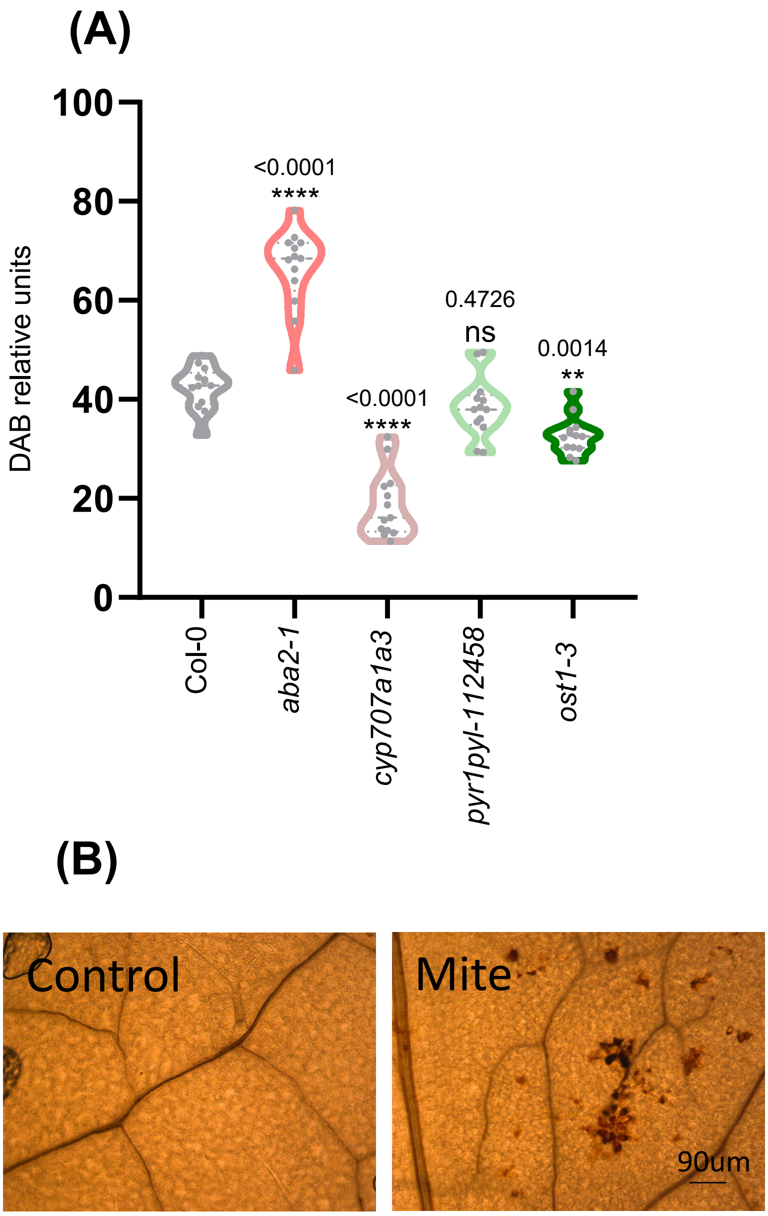


**Figure S5. Redox status in the five Arabidopsis genotypes.** (A) Accumulation of $H_2O_2$ in detached leaves after 24 h of mite infestation, expressed as DAB units. Asterisks and numbers indicate significant differences compared to Col-0 genotype (B) Pictures showing the $H_2O_2$ deposits during *T. urticae* feeding.

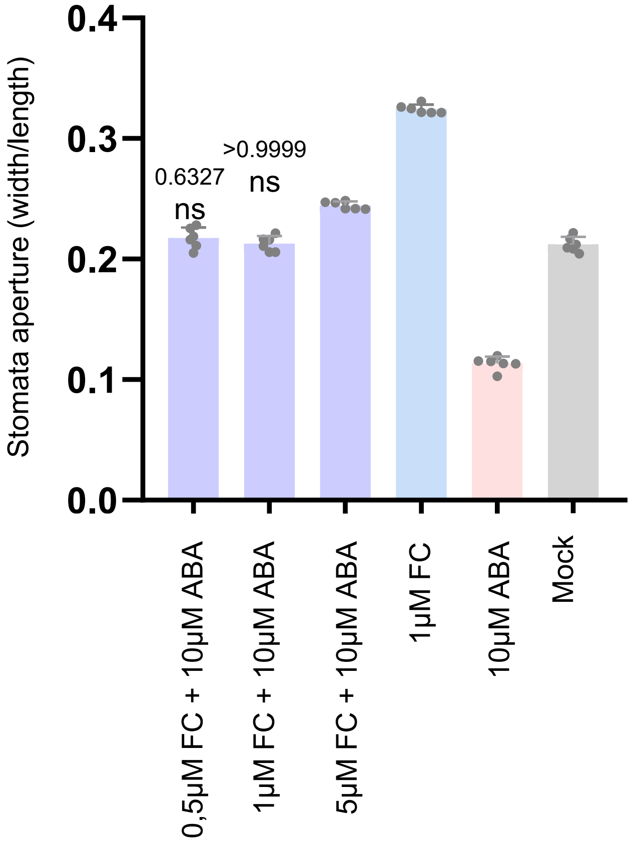


**Figure S6. Effects of different ratios Fussicoccin/ABA on stomata aperture in Arabidopsis Col-0 plants.** Stomata aperture of Arabidopsis plants pre-treated with 10 µM ABA, 1 µM of fusicoccin (FC), or a combination of 10 µM ABA plus different concentrations (0.5, 1.0, and 5.0 µM) of FC. Asterisks and numbers indicate significant differences compare to mock treatment**.**

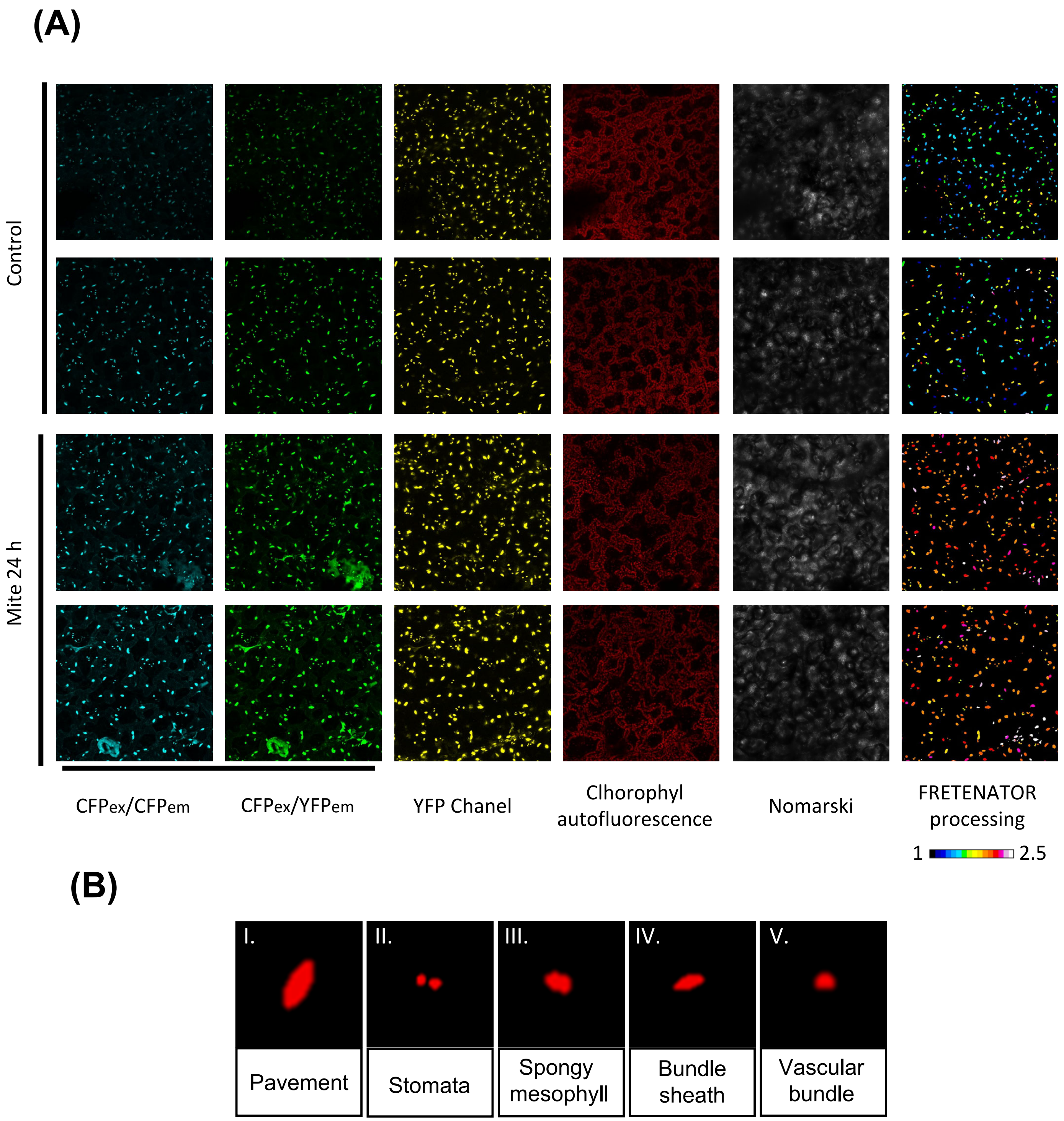


**Figure S7. FRET-images processing.** (A) Raw Confocal images After 24 h of mite infestation (B) FretCellType nucleus determination based on their shape. I. Pavement nuclei. II. Stomata nuclei III. Spongy mesophyll nuclei, IV. Bundle sheat nuclei, V. Vascular bundle nuclei. (A) Raw Confocal images After 24 h of mite infestation
